# Recent origin of Neotropical orchids in the world’s richest plant biodiversity hotspot

**DOI:** 10.1101/106302

**Authors:** Oscar Alejandro Pérez-Escobar, Guillaume Chomicki, Fabien L. Condamine, Adam P. Karremans, Diego Bogarín, Nicholas J. Matzke, Daniele Silvestro, Alexandre Antonelli

**Author notes:** These authors contributed equally to this study (Oscar Alejandro Pérez-Escobar, Guillaume Chomicki). Corresponding authors: Oscar Alejandro Pérez-Escobar.

## Abstract

• The Andean mountains of South America are the most species-rich biodiversity hotspot worldwide with about 15% of the world’s plant species, in only 1% of the world’s land surface. Orchids are a key element of the Andean flora, and one of the most prominent components of the Neotropical epiphyte diversity, yet very little is known about their origin and diversification.

• We address this knowledge gap by inferring the biogeographical history and evolutionary dynamics of the two largest Neotropical orchid groups (Cymbidieae and Pleurothallidinae), using two unparalleled, densely-sampled orchid phylogenies (including 400+ newly generated DNA sequences), comparative phylogenetic methods, geological and biological datasets.

• We find that the majority of Andean orchid lineages only originated in the last 15 million years. Most Andean lineages are derived from lowland Amazonian ancestors, with additional contributions from Central America and the Antilles. Species diversification is correlated with Andean orogeny, and multiple migrations and re-colonizations across the Andes indicate that mountains do not constrain orchid dispersal over long timescales.

• Our study sheds new light on the timing and geography of a major Neotropical radiation, and suggests that mountain uplift promotes species diversification across all elevational zones.

## Introduction

Species richness is unevenly distributed in time (Simpson, 1953), space (Willis, 1922), and across the Tree of Life (Vargas & Zardoya, 2014). Understanding the processes underlying current patterns in species richness and distribution constitutes therefore a major scientific challenge. The Andean mountains of South America contain about 15% of the world’s plant species, in only 1% of the world’s land surface – resulting in the most species-rich biodiversity hotspot worldwide (Myers *et al*., 2000). A large proportion of this diversity is found in high-altitude grasslands, and is suggested to have resulted from recent rapid speciation events (Hughes & Eastwood, 2006; Hughes & Atchison, 2015). By contrast, Andean seasonally dry forests experienced much slower diversification and have older origins (Pennington *et al*., 2010), suggesting contrasted macro-evolutionary histories within the Andean biodiversity hotspot (Valencia *et al*., 1994; Pennington *et al*., 2010; ter Steege *et al*., 2013).

In a seminal paper, (Gentry, 1982) postulated that mountain uplift was a major trigger of Andean mega-diversity, although he posited that this might have occurred indirectly via biotic interactions. A pivotal result of Gentry’s floristic analyses was the discovery of two patterns of plant distribution in the Neotropics: “Amazonian-centered” and “Andean-centered” taxa (Gentry, 1982). Amazonian-centered taxa consist mostly of canopy trees and lianas, while Andean-centered taxa are almost exclusively epiphytes and shrubs (Gentry, 1982). The latter occur mostly in the Northern Andes, with secondary centres in the Brazilian coastal mountains and Central America, together accounting for about 33% of all Neotropical plants (Gentry, 1982) and thus largely contributing the world’s most species-rich biodiversity hotspot, the tropical Andes (Myers *et al*., 2000). Contrasting with dominant views at the time, Gentry (1982) hypothesized that the Andean-centered flora resulted from “recent, very dynamic speciation”.

Orchids are one of the most characteristic and diverse components of the Andean flora (Gentry & Dodson, 1987; Krömer & Gradstein, 2003; Richter *et al*., 2009; Parra-Sánchez *et al*., 2016). They often make up to 30 to 50% of the total epiphytic species number reported along Northern Andes (Kreft *et al*., 2004; Küper *et al*., 2004), and epiphytic orchids account for 69% of all vascular epiphytes worldwide (Zotz & Winkler, 2013). Neotropical epiphytic orchids are generally characterized by narrowly restricted populations with small numbers of individuals (Tremblay & Ackerman, 2001; Jost, 2004; Crain & Tremblay, 2012; Pandey *et al*., 2013). Despite the ecological importance and prominence of epiphytic orchids (and of epiphyte diversity overall) in the Andean flora, their origin and diversification have not been explicitly studied due to the difficulties in generating densely sampled and strongly supported phylogenies.

We address this issue by studying the evolutionary history of the two largest Neotropical orchid clades, namely Cymbidieae and Pleurothallidinae. The Cymbidieae comprise over 3,700 species, 90% of which occur in the Neotropics (remaining species occur in tropical Africa and Australasia). Cymbidieae includes 12 subtribes, of which four are the most speciose and Andean-dwelling subclades (i.e. Maxillariinae, Oncidiinae, Stanhopeinae and Zygopetaliinae; Pridgeon *et al*., 2009). Pleurothallidinae comprise 44 genera and 5100 exclusively Neotropical species (Karremans, 2016) distributed mostly in highlands of the Northern Andes and Central America. Together, they are the most representative elements of the Andean orchid flora (Pridgeon *et al*., 2009; Kolanowska, 2014), and the make up most of their species richness. In addition, these lineages have evolved a rich array of pollination syndromes and sexual systems (Gerlach & Schill, 1991; Pérez-Escobar *et al*., 2016b) that have long fascinated botanists and naturalists (Lindley, 1843; Darwin, 1877). This is particularly true for Cymbidieae, in which up to seven pollination syndromes have been recorded (van der Cingel, 2001; Pridgeon *et al*., 2009), ranging from species exclusively pollinated by male Euglossini bee (Ramirez *et al*., 2011) to those pollinated only by oil bees. Data on pollination ecology of Pleurothallidinae is very scarce, but scattered reports across the clade suggest that they are mostly pollinated by a vast array of Diptera lineages (Blanco & Barboza, 2005; Pupulin *et al*., 2012).

Rapid Andean orogeny could have promoted orchid species richness by creating ecological opportunities such as increasing landscape, mediating local climate change, creating novel habitats, and forming insular environments that affected migrations and allopatric speciation through isolation (Hoorn *et al*., 2013). This effect should have been most accentuated over the last 10 million years (Ma), during which ca 60% of the current elevation of the Andes was achieved (Gregory-Wodzicki, 2000). Diversification studies of Andean-centered clades provide evidence for rapid diversification that temporally matches the Andean surface uplift, notably in the plant genera *Lupinus, Espeletia, Halenia*, *Heliotropium*, and in families Campanulaceae and Annonaceae (von Hagen & Kadereit, 2003; Bell & Donoghue, 2005; Donoghue & Winkworth, 2005; Hughes & Eastwood, 2006; Pirie *et al*., 2006; Antonelli *et al*., 2009b; Luebert *et al*., 2011; Drummond *et al*., 2012; Madriñán *et al*., 2013; Lagomarsino *et al*., 2016). Taken together, these studies suggest that rapid Andean uplift yielded new niches that fostered both adaptive and non-adaptive radiations (Nevado *et al*., 2016). Other Andean groups, such as hummingbirds, diversified mostly prior to Andean uplift (McGuire *et al*., 2014), or after it had attained most of its currently height (Smith *et al*., 2014).

We address the impact of the Andean uplift on the diversity and distribution of orchids by inferring the dynamics of speciation, extinction, and migration while simultaneously incorporating surface uplift of the two largest Andean Neotropical orchid clades Cymbidieae and Pleurothallidinae. We rely on model-based inference methods in historical biogeography, ancestral area and character estimation approaches, and a series of diversification analyses to investigate the following questions: (*i*) Where do Andean orchids come from? (*ii*) Is there evidence for the Andes acting as a dispersal barrier for epiphytic lowland taxa? (*iii*) Did the Andean uplift enhance orchid diversification, and if so was this effect evident on species at all or just the highest elevations? (*iv*) Is Andean diversity derived from pre-adapted lineages or rather descendant of lowland migrants? In addition, we use the limited available data to evaluate whether shifts in pollination syndromes are associated to changes in diversification rates.

Our results support Gentry’s (Gentry, 1982) prediction that Andean-centered groups result from recent rapid speciation, suggesting that Andean orogeny provided opportunities for rapid orchid species diversification in the world’s premier plant biodiversity hotspot. Such diversity is derived from lowland lineages but more rarely from migrants already pre-adapted to cool environments, a more frequent situation documented from other mountain environments (Merckx *et al*., 2015).

## Material and Methods

### Taxon sampling, DNA sequencing and phylogenetic analysis

To generate solid phylogenies of the tribe Cymbidieae and subtribe Pleurothallidianae, we newly generated a total of 420 sequences of the nuclear ribosomal internal transcribed spacer (ITS), and a ~1500 bp fragment of the gene *ycf*1 of under-represented lineages of key biogeographical importance. DNA amplification, PCR product purification and sequencing were conducted as previously described (Pérez-Escobar *et al*., 2016b). Voucher information and GenBank accession numbers are provided in Tables S1 and S2.

We merged our novel dataset with previously generated data from the studies of (Blanco *et al*., 2007), (Whitten *et al*., 2014), and (Ramirez *et al*., 2011), using the R-package MEGAPTERA v.2 (Heibl, 2014). We retrieved 3541 sequences of nuclear (ITS) and plastid (*mat*K, *trn*L-F region, *psb*A, *ycf*1). We selected outgroup taxa representing the old and new world subtribes Polystachyinae, Aeridinae and Laeliinae. Trees were rooted on *Calypso bulbosa* (for Cymbidieae) and *Arpophyllum giganteum* (for Pleurothallidinae) following Whitten *et al*. (2014).

Poorly aligned positions were excluded from the alignments using GBLOCKS v.0.9 (Talavera & Castresana, 2007). To statistically detect potential incongruences between plastid and nuclear DNA phylogenies, we used the tool Procrustes Approach to Cophylogeny (PACo) (Balbuena *et al*., 2013; Pérez-Escobar *et al*., 2016a). Maximum-Likelihood (ML) tree inference was performed using RAxML-HPC v.8.0 (Stamatakis, 2014), under the GTR+G substitution model with four gamma categories (best model for both datasets as inferred via AIC in jModelTest v.2.1.6 (Darriba *et al*., 2012), with 1000 bootstrap replicates and data partitioning by genome compartment. All phylogenetic and dating analyses were performed in the CIPRES Science Gateway computing facility (Miller *et al*., 2015).

### Molecular clock dating

A few unambiguous orchid macrofossils are available for Orchidaceae (*Dendrobium winikaphyllum*, *Earina fouldenensis*, *Meliorchis* caribea (Ramírez *et al*., 2007; Conran *et al*., 2009), but these are assigned to lineages very distantly related to our groups of interest. Using distant outgroups to calibrate our Cymbideae and Pleurothallidinae phylogenies would have created extensive sampling heterogeneities, which can result in spurious results (Drummond & Bouckaert, 2014). Thus, we had to rely on secondary calibrations. In order to obtain the best secondary calibration points possible, we first generated an Orchidaceae-wide fossil-calibrated phylogeny, sampling 316 orchid species sampled as evenly as possible along the tree. Detailed settings and fossil calibrations used to generate an Orchidaceae-wide phylogeny are provided in the extended Material and Methods of Appendix S1.

Secondary calibration points were obtained from our Orchidaceae-wide dated phylogeny, and the MRCA of Cymbidieae + Vandeae was dated to 34 ±7 Ma, 95% credible interval, whereas that of Pleurothallidinae + Laelinae was estimated to 20±7 Ma. We therefore used a normal prior (with values of mean = 34, stdev = 4 for Cymbidieae; mean = 20, stdev = 3 for Pleurothallidinae, to reflect the 95% CI from our fossil-calibrated tree) to calibrate our trees using this secondary constraint, which was designed to reflect the uncertainty previously estimated for the root node of Cymbidieae and Pleurothallidinae.

### Ancestral range estimation

Species ranges were coded from the literature (Pridgeon *et al*., 2009) and from herbarium specimens through a survey of virtual collections and loans of several herbaria (AMES, COL, F, MO, SEL, US, M) as well as the GBIF repository. To query GBIF database, we relied on the function *occ* of the R-package SPOCC (Scott *et al*., 2015). A total of 19486 distribution records were compiled for the Cymbidieae, and 9042 records for the Pleurothallidinae. Protocols for distribution maps and species richness pattern analyses are detailed in Appendix S1.

Distribution maps for Cymbidieae and Pleurothallidinae (summarized in Figs. S1, S2) and extant distribution patterns identified for other plant lineages (e.g. Rubiaceae [Antonelli *et al*., 2009b]) allowed the identification of 10 main distribution areas (see the inset in Fig. 1 and 2). Species were assigned to one of these regions: (*i*) Central America (comprising southern Florida to Panama); (*ii*) West Indies; (*iii*) Northern Andes (mountain ranges from elevations higher than 500 m in Colombia and Venezuela); (*iv*) Central Andes (from Peru to the Tropic of Capricorn, from elevations higher than 500 m); (*v*) Amazonia, including lowlands and montane forest below 500 m in Colombia, Ecuador, Peru, Brazil, Venezuela, Guyana, Suriname and French Guiana; (*vi*) The Guiana shield, including elevations higher than 500 m in north-eastern South America (Brazil, Guyana, Suriname and Venezuela); (*vii*) Southeastern South America, including the Brazilian shield but also lowlands in eastern Brazil and Northern Argentina; (*viii*) Chocó (comprises lowlands below 500 m of the western Andes in Colombia, Ecuador); (*ix*) Africa and (*x*) Australasia. To infer the ancestral range of all examined lineages in Cymbidieae and Pleurothallidinae, we used the R package BioGeoBEARS v.0.2.1 (Matzke, 2013, 2014). In addition, in order to estimate the number of migrations, dispersals, extinctions and sympatric speciation events from our phylogeny, we used Biogeographical Stochastic Mapping (BSM) (Matzke, 2014) under the best-fit model, as implemented in BioGeoBEARS (for detailed settings see Appendix S1).

**Figure 1.**
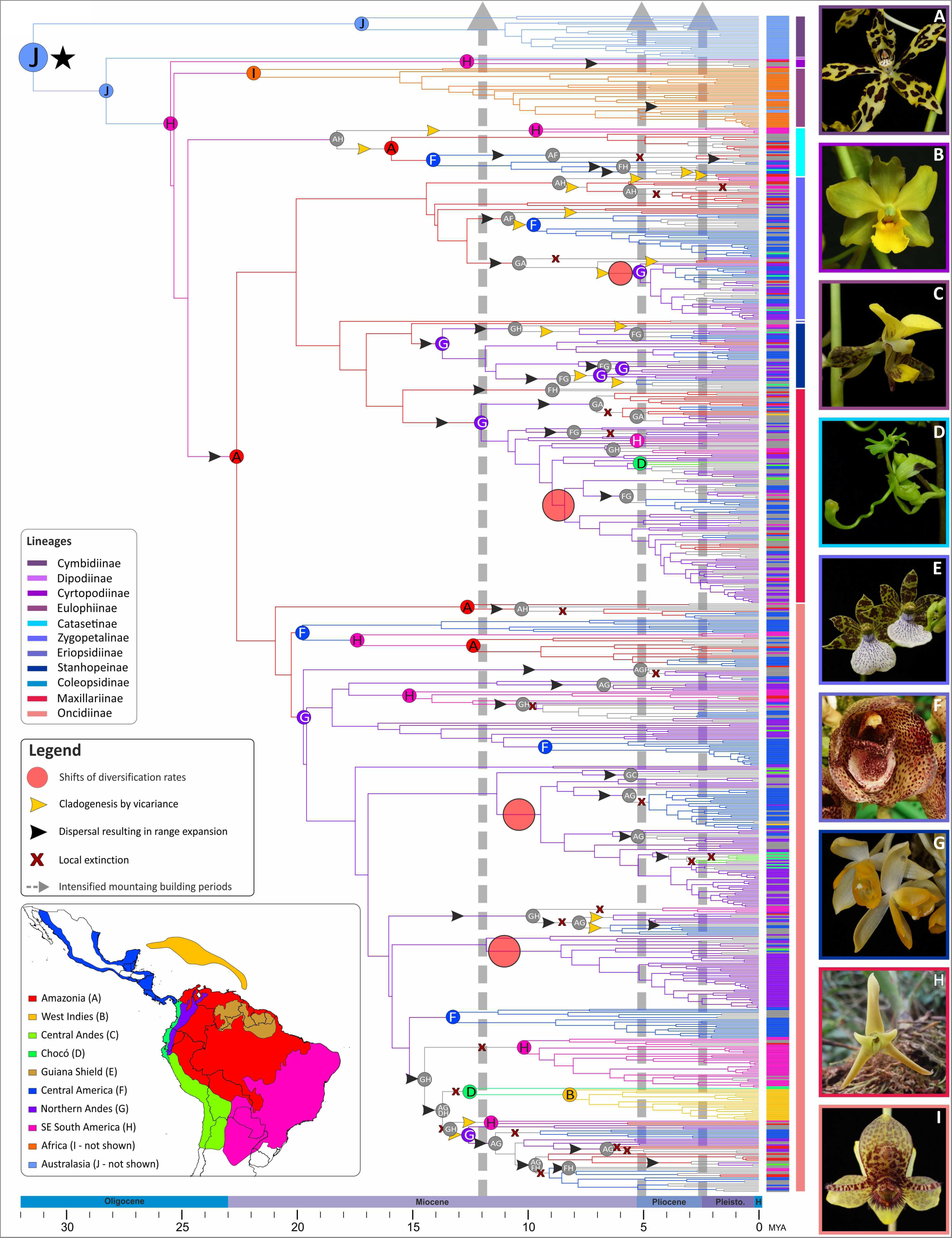
Biogeographic history of Cymbidieae orchids. Letters on colored circles at nodes indicate the estimated ancestral area with the highest probability as inferred by BioGeoBEARS. Branches are color-coded following the reconstructed area of their corresponding node, and geographical ranges of every taxon are shown as vertical bars in front of the terminals. The black start indicates the MRCA of Cymbidieae. Gray arrows show the periods of accelerated Andean uplift (Gregory-Wodzicki, 2000). Changes on shifts of diversification rates are shown as red pale circles on branches. Range expansions, local extinctions and cladogenetic events via vicariance are indicated on branches with black, yellow arrows and red crosses, respectively. Sub-tribe members of Cymbidieae are color-coded. Right panels show selected representatives of (a) Cymbidiinae (*Grammatophyllum measuresianum*); (b) Cyrtopodiinae (*Cyrtopodium polyphyllum*; photo by Luiz Varella); (c) Eulophiinae (*Eulophia streptopetala*); (d)Catasetinae (*Cycnoches egertonianum*); (e) Zygopetaliinae (*Zygopetalum* aff. *brachypetalum*). (f) Coeliopsidiinae (*Peristeria cerina*); (g) Stanhopeinae(*Sievenkingia* sp.); (h) Maxillariinae (*Cryptocentrum* sp.); (i) Oncidiinae (*Trichoceros* sp.). Photos (except b): O. Pérez. (Inset) Coded areas for biogeographical analysis. Political divisions obtained from DIVA-GIS (http://www.diva-gis.org/gdata).

**Figure 2.**
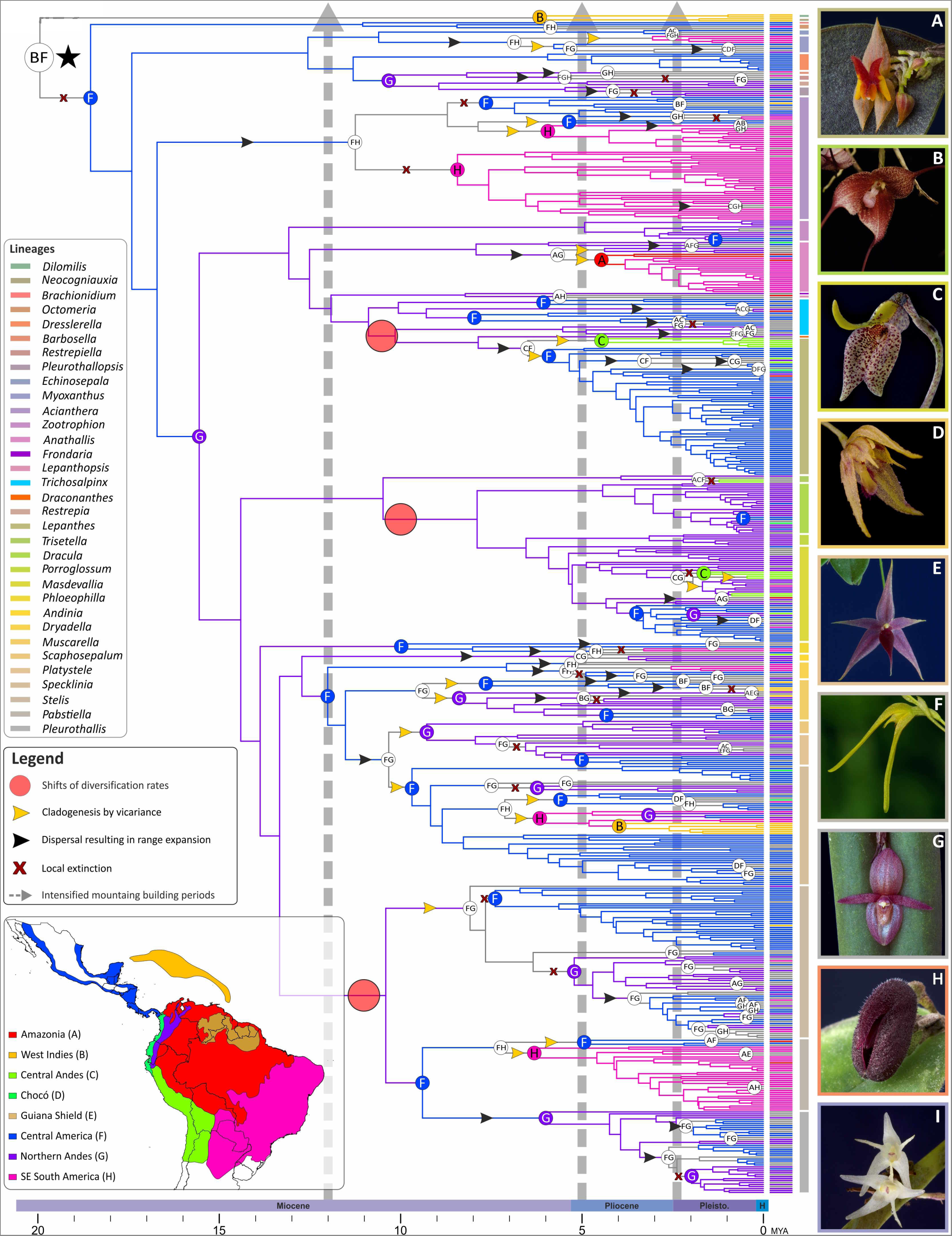
Biogeographic history of Pleurothallidinae orchids. Letters on colored circles at nodes indicate the estimated ancestral area with the highest probability as inferred by BioGeoBEARS. Branches are color-coded following the reconstructed area of their corresponding node, and geographical ranges of every taxon are shown as vertical bars in front of the terminals. The black start indicates the MRCA of Pleurothallidinae. Gray arrows show the periods of accelerated Andean uplift (Gregory-Wodzicki, 2000). Changes on shifts of diversification rates are shown as red pale circles on branches. Range expansions, local extinctions and cladogenetic events via vicariance are indicated on branches with black, yellow arrows and red crosses, respectively. Generic members of Pleurothallidinae are color-coded. Right panels show selected representatives of *(a) Lepanthes* (*Lepanthes* sp.); (b) *Dracula* (*D. astuta*); (c) *Masdevallia* (*M. utricularia*); (d) *Muscarella* (*M. exesilabia*); (e) Platystele (*P. porquinqua*); (f) *Pabstiella* (*P. ephemera*); (h) *Pleurothallis* (*P. adventurae*); (i) *Myoxanthus* (*M. colothrix*). Photos: A. Karremans, D. Bogarín and O. Pérez. (Inset) Coded areas for biogeographical analysis. Political divisions obtained from DIVA-GIS (http://www.diva-gis.org/gdata).

### Rates of species diversification

To infer the diversification dynamics of the Cymbidieae and Pleurothallidinae, we first used a time-dependent model implemented in BAMM v.2.5.0 (Rabosky, 2014) to estimate rates of extinction and speciation across the phylogenies. Incomplete taxon sampling was accounted for by assigning a sampling fraction of 25% of the extant orchid diversity of Cymbidieae, and 13% of Pleurothallidinae (sampling fractions of every genus sampled was incorporated according to [Chase *et al*., 2015]). We performed three runs with 1 million MCMC generations, sampling parameters every 10,000 generations. Diversification rates and rate shift configurations were plotted using the R package BAMMtools (Rabosky *et al*., 2014). We checked the convergence of the runs by plotting the log-likelihood across MCMC generations sampled in the “mcmc_out” file. To evaluate the best model generated by BAMM (compared with a null *M_0_* model with no diversification rates shifts), we relied on Bayes Factors calculated with the *ComputeBayesFactor* function of BAMMtools. We examined the 95% credible set of macroevolutionary shift configuration using the BAMMtools function *credibleShiftSet*. Settings for the BAMM cross validation analysis carried in RPANDA (Morlon *et al*., 2016) are provided in Appendix S1.

### Geographic state-dependent analyses

We used GeoSSE (Goldberg *et al*., 2011), an extension of the BiSSE model that allows lineages to occur simultaneously in two areas and to test whether one area has overall higher speciation rates, as implemented in the R package *diversitree* v.0.9-7 (Fitzjohn, 2012). To test whether lineages restricted to the Northern Andes (“A”) had higher diversification rates than lineages absent from the Northern Andes (collectively called “B” here), we used Bayesian MCMC GeoSSE analyses of 1 million generations on the maximum clade credibility tree from BEAST (in the particular case of Cymbidieae, only Neotropical representatives were included). Implemented models in GeoSSE and settings of tailored simulation to account for Type I error bioases in GeoSSE are provided in Appendix S1.

### Mapping speciation rate in the Neotropics

Based on the speciation and extinction rates inferred for orchid lineages, and their geographic occurrence, it is possible to identify important areas of diversification as plotted on a heat map (Condamine & Kergoat, 2013). For this purpose, we designed a novel method that consists on retrieving speciation (lambda) rates from BAMM analyses using the function *GetTipsRates* in BAMMtools v.2.1 (Rabosky *et al*., 2014) and to link them to species occurrences. Rates were further associated to known distribution records of Cymbidieae and Pleurothallidinae and interpolated to a polygon representing the currently known distribution of Cymbidieae and Pleurothallidinae species, using the Inverse Distance Weight method implemented in the software ArcMap v.9.3 (Esri). To account for geographical sampling biases from herbarium records, we randomly sampled from 0.5x0.5-degree grid cells herbarium occurrences using the R package RASTER (Hijmans & Elith, 2016) so that a single occurrence per grid cell was kept.

### Paleoelevation-dependent diversification

We tested the effect of past environmental change on the diversification of Cymbidieae and Pleurothallidinae using birth-death models that allow speciation and extinction rates to vary according to a quantitative, time-dependent environmental variable (Condamine *et al*., 2013), here the paleo-elevation of the northern Andes (Hoorn *et al*., 2010). The R-package PSPLINE (Ramsey & Ripley, 2010) was used to interpolate a smooth line for Andean paleo-elevation. This smooth line was sampled during each birth-death modeling process to give the value of the paleoelevation variable at each time point. Speciation and extinction rates were then estimated as a function of these values along the time-calibrated phylogenies, according to the parameters of each model. The paleo-environmental-dependent model is implemented in the R-package RPANDA v.1.1 (Morlon *et al*., 2016). Implemented models in RPANDA are provided in Appendix S1.

### Ancestral character state estimation

To account for potential biotic variables as drivers of Neotropical orchid diversification such as shifts on pollination syndromes (Givnish *et al*., 2015), we compiled information on pollination syndromes of Cymbidieae from the literature (van der Cingel, 2001; Singer, 2002; Pansarin *et al*., 2009; Pridgeon *et al*., 2009; Gerlach, 2011; Ramirez *et al*., 2011), and consulted experts on specific groups (see *Acknowledgements*). Due to a dearth of detailed information on pollination ecology (i.e. ~6% of taxa sampled), we followed a generalist coding approach, and seven pollination syndromes, (i.e. bee, bird, butterfly, lepidopteran, fly, wasp and self-pollination) were coded. To account for missing information on pollination syndromes, we assigned equal probabilities to all character states to taxa with unknown pollination syndrome. To estimate ancestral elevation ranges in Pleurothallidinae and Cymbidieae, we obtained absolute elevation values from herbarium records for every taxon sampled in our phylogenies. We obtained a mean of five values per taxa sampled, and we coded mean elevation values as a continuous character. Detailed settings for Ancestral Character State (ACS) of altitude and pollination syndromes are provided in Appendix S1.

## Results

### Phylogenetics, age and biogeography of Andean orchids

Analyses of phylogenetic incongruence detection identified 259 and 125 potential conflicting tips in Cymbidieae and Pleurothallidianae, respectively (Fig. S27, 28), all of which clustered in weakly to moderately supported clades (<75% LBS), or in clades with extremely long branches. In the absence of supported phylogenetic conflicts, nuclear and chloroplast partitions of Cymbideae and Pleurothallidinae were concatenated. For the Cymbidieae, our molecular dataset consisted of 6.6 kb DNA (five markers) for 816 species, and yielded the first strongly supported phylogeny of the tribe (Fig. S3). The Pleurothallidinae dataset was composed of 2.4 kb DNA (two markers) and 684 terminals, including in total 420 newly generated sequences (Fig. S4). Both orchid phylogenies are strongly supported at most important nodes, with 618 nodes (76%) with a maximum likelihood bootstrap support (BS) > 75% for the Cymbidieae, and 321 nodes (47%) for the Pleurothallinae (Figs. S3, S4).

Ages obtained on our wide-orchid dated phylogeny were very similar from other recent orchid dating studies (Chomicki *et al*., 2015; Givnish *et al*., 2015). A chronogram for the orchid family showing absolute ages and 95% confidence intervals for every node is provided in Fig. S5. Absolute ages obtained for Cymbidieae and Pleurothallidinae chronograms are also in agreement with previously published dated phylogenies (e.g. Ramirez *et al*., 2011; Chomicki *et al*., 2015; Givnish *et al*., 2016). Divergence time estimates and 95% credible intervals inferred for all nodes of Cymbidieae and Pleurothallidinae chronograms are shown in Fig. S6 and S7.

Our dating and biogeographic analyses identified DEC + J as best fitting model for both Cymbidieae and Pleurothallidinae (Table S3, S4). Under this model, an Australasian origin of the Cymbidieae around the Eocene/Oligocene boundary (34±8 Ma) was inferred (Fig. 1, Figs. S6, S8). We inferred a late Oligocene dispersal from Australasia to South America following the estimation of southern South America as ancestral area of *Cyrtopodium* and the rest of the Cymbidieae (Fig. 1; Fig. S8). Such dispersal corresponds to the final break-up of Gondwana (split between Antarctica and South America at Drake Passage). From the late Oligocene to the early Miocene, our analyses indicate dispersal from East to West in the Neotropics. The Northern Andean region was reached three times from Amazonia by the most recent common ancestor (MRCA) of Oncidiinae ca. ~19±5 Ma, Maxillariinae ca. 11±5 Ma, and Stanhopeinae ca. 13±4 Ma.

Ancestral state estimations of mean altitude further show that the MRCAs of Cymbidieae was likely adapted to lowland environments (ancestral elevation value of 900 m; Fig. S9, S10). The MRCAs of Amazonian migrants that reached the Andes (i.e. Maxillariinae, Oncidiinae and Stanhopeinae) were also adapted to lowland habitats (mean elevation values of ~1300, 1200, and 900 m, respectively; Fig S9, S10). Strikingly, Oncidiinae and Maxillarinae are the species richest lineages in Cymbidieae (1,584 and 819 species, respectively; Chase *et al*., 2015), and they are derived from lowland Amazonian migrators. Stanhopeinae subsequently dispersed to several other Neotropical regions, notably Central America (Fig. 1, Fig. S8).

Different from the Cymbidieae, we infer an origin of Pleurothallinae in Central America or the West Indies in the early Miocene, followed by a migration to the Northern Andes *ca.* 16±5 Ma (Fig. 2, Fig. S7, S11), prior to the main uplift periods but at a time frame when the Northern Andes had already peaked mean elevations of ~1500 m. However, the majority of early divergent Pleurothallidinae and their sister groups are from the Antilles, and thus the inference of Central America as the ancestral area of Pleurothallidinae most likely reflects our inability to sample extensively the early divernging Antillean lineages. As inferred by ancestral state estimations, the MRCA of Pleurothallidinae was adapted to montane habitats (mean elevation of ~1200 m), and all Pleurothallidinae migrants to the Northern Andes were likely adapted to montane–cloud forest environments (mean elevation of ~1500 m; Fig. S12-S13). Biogeographical stochastic mapping indicates that in-situ speciation was the dominant biogeographic process in both clades, while processes of range expansion (dispersal and vicariance) and range contraction (subset speciation) were scarcer and relatively evenly distributed across lineages (Fig. 1–2; Fig. S14-S15).

### Diversification of Andean orchids

The diversification analyses performed with BAMM strongly rejected a constant-rate model (Bayes factor=151.3, Table S5), and instead identified four rate shifts during the evolutionary history of Cymbidieae (Fig. 3B; Figs. S16, S17). The best model configuration identified four shifts in speciation rate in the most speciose Cymbidieae lineages: one in Maxillariinae, one in Zygopetalinae, and two in Oncidiinae. We further identified three rate shifts in the Pleurothallinae (Table. S6), in the MRCAs of *Lepanthes* + *Lepanthopsis*, *Dracula* + *Porroglossum* + *Masdevallia*, and *Stelis* + *Pabstiella* + *Pleurothallis* (Fig. 4B; Fig S18, S19). All shifts in diversification rates in Cymbidieae and Pleurothallidiane were further confirmed using the RPANDA method (Fig. S20, 21; Tables S7, S8).

**Figure 3.**
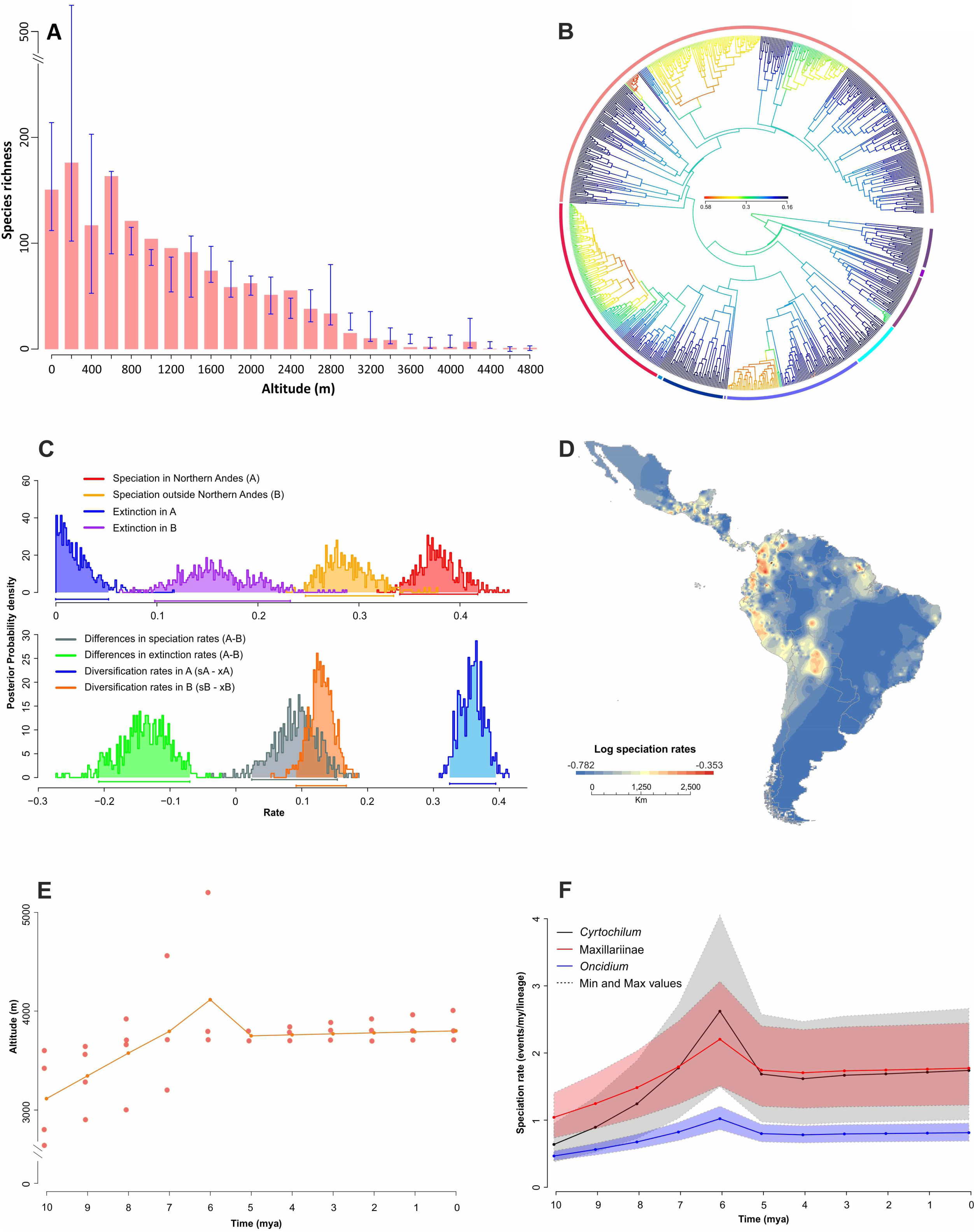
Diversification of the Cymbidieae. (A) Richness through elevation plot for 55% (>20,000 herbarium records) of the ca. 4000 Cymbidieae species. (B) Speciation rate plot (phylorate) showing the best configuration shift identified by BAMM. (C) Density probability plots of speciation, extinction and net diversification rates per area identified by GeoSSE. Area “A” refers to species restricted to the Northern Andes; Area “B” refers to species occurring in all areas except Northern Andes. (D) Speciation rate map estimated from BAMM (Materials and Methods). (E) Average paleoelevation of the central and northern Andes. (F) Paleoelevation-dependent models applied to the four clades detected by BAMM to have significantly higher diversification rates than others. Lineages in (B) are color coded in the same way as shown in Figure 1.

**Figure 4.**
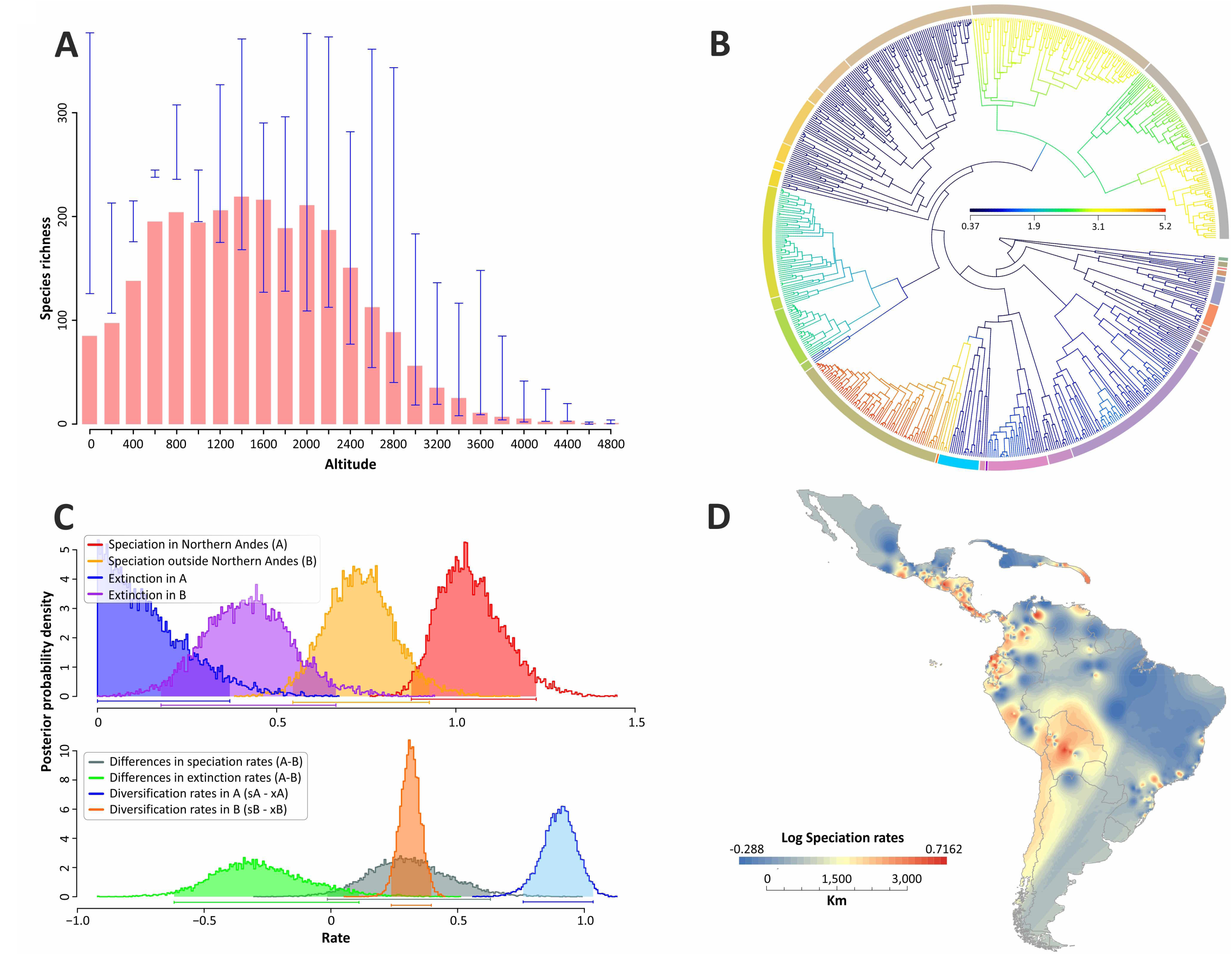
Diversification of the Pleurothallidinae. (A) Richness through elevation plot for 50% (>9000 herbarium records) of the ca. 5000 Pleurothallidinae species. (B)Speciation rate plot (phylorate) showing the best configuration shift identified by BAMM. (C) Density probability plots of speciation, extinction and net diversification rates per area identified by GeoSSE. Area “A” refers to species restricted to the Northern Andes; Area “B” refers to species occurring in all areas except Northern Andes. (D) Speciation rate map estimated from BAMM (Materials and Methods). Lineages in (B) are color coded in the same way as shown in Figure 2.

Interestingly, the diversification rate shifts are all located at the origin of Northern Andean clades and temporally match with periods of accelerated Andean uplift in this region (Cymbidieae, Fig. 1), or within a clade that already inhabited the Northern Andes (e.g. Pleurothallinae, Fig. 2). To further explore this apparent correlation with either accelerated Andean uplift or presence in the Northern Andes and fast diversification, we used a trait-dependent approach (GeoSSE) that estimates region-dependent speciation rates. Here, a model with free rates fitted best our Cymbidieae and Pleurothallidinae dataset (Table S9), with differences in speciation (sA – sB) and diversification (dA - dB) rates highly if not maximally supported (0.99 and 1 Bayesian Posterior Probabilities, respectively). GeoSSE analyses further indicated that speciation rates in Northern Andes are consistently higher than in any other biogeographical region (Fig 3C, 4C) in both Cymbidieae and Pleurothallidinae datasets. We evaluated and confirmed the robustness of these results through extensive data simulations (Fig. S21). Here, the distribution delta AIC obtained from AIC values from real data and reshuffled data analyses was centered towards -20 and far away from 0 (i.e. AIC values obtainedof real data set).

We developed a novel method to generate a ‘speciation rate map’ using inferred speciation rates for each orchid lineage and georeferenced species occurrences (see *Materials and Methods*). Our speciation rate maps are in agreement with GeoSSE results, and we confirmed that speciation rates in the Northern Andes were significantly higher than those in any other region (Figs. 3C, 4C). This is in agreement with a recent study with more limited taxon sampling for the two clades focused here (Givnish *et al*., 2015). The speciation rate map (*Materials and Methods*) further demonstrates that fastest speciation took place in the Northern Andes region, and reveals secondary speciation hotspots in the Central Andes, the Guiana Shield, and Central America (Fig. 3D, 4D). These secondary hotspots are occupied by species derived from the four highly diversifying Northern Andean Cymbidieae clades (Fig. S23), suggesting that the Andes acted as a major source of new lineages to the rest of the continent – thus greatly increasing Neotropical orchid diversity. This is particularly true for the Pleurothallidinae, where we identified multiple migrations from the Northern Andes of montane adapted lineages to Central America (Fig 2; Fig. S24). Interestingly, we also found a strong geographic correlation between current species richness and diversification (Figs. 3D, 4D, S25, S26), suggesting that recent *in-situ* speciation was the main process for species accumulation in the Neotropics.

While these results suggest an impact of the Andean uplift on species diversification, they do not explicitly account for biotic interactions, landscape and climatic changes through time. We therefore assessed the fit of a model that explicitly integrates paleo-elevation in diversification rate analyses (see *Materials and Methods*). In three of the four Cymbidieae clades where BAMM inferred a speciation rate shift, the paleo-elevation-dependent model inferred a continuous speciation increase from 10 to 6 Ma (Fig. 3E-F, Table S10). In contrast, no positive correlation with paleo-elevation and diversification could be detected for Pleurothallidinae (Table S11). Moreover, our ancestral character estimation of pollination syndromes in Cymbidieae suggests that the MRCA of Cymbidieae was bee pollinated (Fig. S29). Nine shifts of syndromes were identified along the evolutionary history of Cymbidieae, always derived from bee pollination. No reversals from other syndromes towards bee pollination were recovered (Fig. S29).

## Discussion

### Andean orchids are mostly derived from lowland Amazonian migrants

Our ancestral area estimations show that Andean orchid flora is derived primarily from Amazonian lowland taxa (i.e. MRCAs of Oncidiinae, Maxillariinae and Stanhopeinae, the most speciose lineages in Cymbidieae), but to a lesser extent also from cool pre-adapted lineages (MRCA of most extant Andean centred Pleurothallid taxa). Previous research has revealed that mountain flora origin is strongly influenced by the immigration of cool pre-adapted lineages (Hughes & Eastwood, 2006; Merckx *et al*., 2015; Uribe-Convers & Tank, 2015), and that contributions from lowland adapted lineages is rather rare. Our study points to the key role of Amazonia for the origin of Andean orchid diversity, and also reveals an ancient biological connectivity between this region and the Northern Andes.

### The Andes did not constrain orchid dispersal

The recurrent migration back and forth through the Andes, even during the period of highest paleo-elevation, is also a central result from our study. The colonization of the Northern Andes by some clades of Cymbidieae matches in time with accelerated surface uplift (Fig. 1, Fig. S8), and reflects the Miocene biotic connectivity between the Andes and Amazonia previously suggested for plants (Antonelli *et al*., 2009a), Poison dart frogs (Santos *et al*., 2009), and birds (Brumfield & Edwards, 2007), among others. This suggests that shifts across elevational zones were rare, similarly to recent results in Mount Kinabalu in Borneo (Merckx *et al*., 2015).

Surprisingly, dispersal events across the Andes did not decrease during accelerated Andean uplift (Fig. 1, 2; Fig. S8, S11), suggesting that the uplift of the Andes did not act as a major dispersal barrier for Cymbidieae and Pleurothallidinae orchids, contrary to findings in other plant groups (e.g.: Annonaceae [Pirie *et al*., 2006], Rubiaceae [Antonelli *et al*., 2009b] or Fabaceae [Pennington *et al*., 2010]. This result likely relates to the biology of orchids, which produce large amounts of dust-like, wind-dispersed seeds allowing for occasional long-distance dispersal (Arditti & Ghani, 2000; Antonelli *et al*., 2009a; Barthlott *et al*., 2014; Givnish *et al*., 2016). Taken together, these findings suggest that the Andes constitute a semi-permeable barrier to biotic dispersal, and that orchids may be more geographically constrained by intrinsic factors such as fungal symbionts and pollinators, which differ among elevational zones (Arroyo *et al*., 1982, 1985; Lugo *et al*., 2008) than by distance.

### Accelerated orchid diversification across elevational zones

Gentry’s hypothesis (Gentry, 1982) of rapid speciation in the Andes was mainly based on the observation of floristic groups (“Andean-centered taxa”) with very speciose genera from the lowlands to mid-elevations in the (mostly Northern) Andes. This matches well the total altitudinal distribution of our respective study groups, with a richness-through-elevation plot for >55% of the 3,700 Cymbidieae species based on over 20,000 records (Fig. 3A; Fig. S1), which reveals that Cymbidieae diversity peaks at low elevations, while Pleurothallidinae (ca. 10,000 records; Fig. S2) peaks at mid-elevation at around 1,500 m (Fig. 4A).

The diversification rate shifts are all located at the origin of Northern Andean clades and temporally match with periods of accelerated Andean uplift in this region (Gregory-Wodzicki, 2000; Hoorn *et al*., 2010) (Cymbidieae, Fig. 1), or within a clade that already inhabited the Northern Andes (e.g. Pleurothallinae, Fig. 2). This is the period with fastest documented rates of Andean uplift in the Northern Andes (i.e. Venezuelan Andes and Northern Andes of Colombia; Hoorn *et al*., 1995; Bermúdez *et al*., 2015). In all three Cymbidieae clades, speciation rates peaked at 6 Ma, a time when the northern Andes reached ca. 4,000 m, their maximum mean paleoelevation (Bermúdez *et al*., 2015). Contrary to Cymbidieae, we found no correlation between Andean uplift and Pleurothallidinae diversification (Table S11), consistent with the earlier colonization of the Northern Andean region. We hypothesize that is due to rapid radiation of migrating cool pre-adapted Pleurothallidinae lineages from Central America into already formed montane environments (Hoorn *et al*., 2010). Similar radiation patterns have been already reported for *Lupinus, Bartsia*, Adoxaceae and Valerianaceae (Donoghue & Sanderson, 2015; Uribe-Convers & Tank, 2015).

Gentry proposed that the main mechanism underlying rapid speciation in the Andes was the evolution of novel plant-insect interactions (Gentry, 1982). The Cymbideae are particularly known among biologists and ecologists because of the rich array of pollination syndromes and sexual systems they have evolved (e.g. sexual and food deceit, food and fragrance reward, dichogamy and environmental sex determination [Gerlach & Schill, 1991; Singer, 2002; Pansarin *et al*., 2009; Gerlach & Pérez-Escobar, 2014]). Our analyses suggest that pollinator syndrome shifts do not match with diversification rates shifts, although our data do not take into account pollinator shifts within given pollinator groups. This is particularly true for bee pollination syndrome, which is widespread in the tribe and likely overarch several transitions from different types of bee pollinator (e.g. oil to Euglossini bees as observed in Catasetinae). More field observations of pollinations are therefore needed to evaluate the relative role of pollinator shifts in contributing to Neotropical orchid diversification.

## Conclusion

Based on two extensively sampled orchid phylogenies combined with statistically robust diversification models, our results reveal that Andean orchid diversification have closely tracked the Andean orogeny. Together with studies in other mega-diverse regions (Verboom *et al*., 2009; Bruyn *et al*., 2014), our results show that rapid recent speciation has moulded this area of exceptional species richness. In addition, our results highlight the crucial role of Amazonian lowlands as well as the Antillean and Central American regions as biotic sources for Andean biodiversity, providing cool pre-adapted lineages that dispersed into the Andes and further diversified *in situ*.

Contrary to general expectation, the rise of the Andes had little effect on restricting orchid biotic dispersal across the Neotropics. This suggests that mountains are semi-permeable barriers to lowland organisms, whose dispersal ability are more likely related to intrinsic traits (e.g. seed size, dispersal mechanism, mutualisms). Although both abiotic and biotic processes are clearly responsible for the exceptional species richness of the world’s premier biodiversity hotspot (Antonelli & Sanmartín, 2011; Hughes *et al*., 2013), our results suggest that geological processes played a central and direct role in the diversification process. Finally, since the highest species richness in Cymbidieae is concentrated in the lowlands and the Pleurothallinae peak at mid-elevation, our study shows that Andean uplift dramatically affected the evolutionary assembly of both lowland and mid-elevation Andean forests, as originally hypothesized by Gentry (1982).

## Author contributions

O.A.P., G.C. and A.A. designed research; O.A.P., A.K., D.B. and G.C. performed research; O.A.P., G.C., F.C., and N.M. analysed data; F.C. and D.S. contributed analytic tools; G.C. and O.P. wrote the paper with contributions from all authors.

## Acknowledgments

We thank M. Whitten, G. Gerlach, and E. Pansarin for plant material; G. Gerlach, M.A. Blanco and G. Salazar for providing information on pollination. C. Bernau for dedicated IT support; A. Mulch for the compilation of surface uplift data from the Andes; L. Lagomarsino for discussion F. Pupulin and B. Gravendeel for their support on the Cymbidieae and Pleurothallidinae research. O.A.P. is supported by Colombian National Science Foundation (COLCIENCIAS) scholarship, G.C. is supported by the German Science Foundation grant (RE 603/20). F.L.C is supported by a Marie Curie grant (BIOMME project, IOF-627684). A.P.K and D.B. were supported by grants of the Alberta Mennega Foundation. N.J.M. was supported by the National Institute for Mathematical and Biological Synthesis, an Institute sponsored by the National Science Foundation through NSF Award #DBI-1300426, with additional support from The University of Tennessee, Knoxville, and is currently supported by Discovery Early Career Researcher Award DE150101773, funded by the Australian Research Council, and by The Australian National University. D.S. is funded by the Swedish Research Council (2015-04748). A.A. is supported by grants from the Swedish Research Council, the European Research Council under the European Union’s Seventh Framework Program (FP/2007-2013, ERC Grant Agreement n. 331024), the Swedish Foundation for Strategic Research and a Wallenberg Academy Fellowship.

## Online Supplementary Material (OSM)

Appendix 1: Extended Materials and Methods

Supplementaryfigures S1-S29

Supplementary tables S1-S11

## References

Antonelli A, Dahlberg CJ, Carlgren KHI, Appelqvist T. 2009a. Pollination of the Lady’s slipper orchid (Cypripedium calceolus) in Scandinavia-taxonomic and conservational aspects. Nordic Journal of Botany 27: 266–273.

Antonelli A, Nylander JAA, Persson C, Sanmartı I. 2009b. Tracing the impact of the Andean uplift on Neotropical plant evolution. 106: 9749–9754.

Antonelli A, Sanmartín I. 2011. Why are there so many plant species in the Neotropics? Taxon 60: 403–414.

Arditti J, Ghani AKA. 2000. Numerical and physical properties of orchid seeds and their biological implications. New Phytologist 145: 367–421.

Arroyo MTK, Armesto JJ, Primack RB. 1985. Community studies in pollination ecology in the high temperate Andes of Central Chile II. Effect of temperature on visitation rates and pollination possibilities. Plant Systematics and Evolution 149: 187–203.

Arroyo MTK, Primack RB, Armesto JJ. 1982. Community studies in pollination ecology in the high temperate Andes of Central Chile I. Pollination mechanisms and altitudinal variation. American Journal of Botany 69: 82–97.

Balbuena JA, Míguez-Lozano R, Blasco-Costa I. 2013. PACo: a novel procrustes application to cophylogenetic analysis. PloS one 8: e61048.

Barthlott W, Große-Veldmann B, Korotkova N. 2014. Orchid seed diversity: a scanning electron microscopy survey. Berlin: Botanic Garden and Botanical Museum Berlin.

Bell CD, Donoghue MJ. 2005. Phylogeny and biogeography of Valerianaceae (Dipsacales) with special reference to the South American valerians. Organisms, Diverisity and Evolution 5: 147–159.

Bermúdez M, Hoorn C, Bernet M, Carrillo E, Beek PA, Van Der, Mora JL, Mehrkian K. 2015. The detrital record of late-Miocene to Pliocene surface uplift and exhumation of theVenezuelan Andes in the Maracaibo and Barinas foreland basins. Basin Research: 10.1111/bre.12154.

Blanco MA, Barboza G. 2005. Pseudocopulatory Pollination in Lepanthes (Orchidaceae: Pleurothallidinae) by Fungus Gnats. Annals of Botany 95: 763–772.

Blanco MA, Carnevali G, Whitten WM, Singer RB, Koehler S, Williams NH, Ojeda I, Neubig KM, Endara L. 2007. Generic realignment in Maxillariinae (Orchidaceae). Lankesteriana 7: 515–537.

Brumfield RT, Edwards SV. 2007. Evolution into and out of the Andes: a Bayesian analysis of historical diversification in Thamnophilus antshrikes. Evolution 61: 346–369.

Bruyn M, Stelbrink B, Morley R, Hall R, Carvalho G, Cannon C, van den Bergh G, Meijaard E, Metcalfe I, Boitani L, et al. 2014. Borneo and Indochina are major evolutionary hotspots for Southeast Asian biodiversity. Systematic Biology 63: 879–901.

Chase MW, Cameron KM, Freudenstein J V., Pridgeon AM, Salazar G, van den Berg C, Schuiteman A. 2015. An updated classification of Orchidaceae. Botanical Journal of the Linnean Society 177: 151–174.

Chomicki G, Bidel LPR, Ming F, Coiro M, Zhang X, Wang Y, Baissac Y, Jay-allemand C, Renner SS. 2015. The velamen protects photosynthetic orchid roots against UV-B damage, and a large dated phylogeny implies multiple gains and losses of this function during the Cenozoic. New Phytologist 205: 1330–1341.

van der Cingel N. 2001. An atlas of orchid pollination: America, Africa, Asia and Australia. Rotterdam: Balkema Publishers.

Condamine FL, Kergoat GJ. 2013. Global biogeographical pattern of swallowtail diversification demonstrates alternative colonization routes in the Northern and Southern Hemispheres.

Conran JG, Bannister JM, Lee DE. 2009. Earliest orchid macrofossils: Early Miocene Dendrobium and Earina (Orchidaceae: Epidendroideae) from New Zealand. American Journal of Botany 96: 466–474.

Crain B, Tremblay R. 2012. Update on the distribution of Lepanthes caritensis, a rare Puerto Rican endemic orchid. Endangered Species Research 18: 89–94.

Darriba D, Taboada GL, Doallo R, Posada D. 2012. jModelTest 2: more models, new heuristics and parallel computing. Nature Methods 9: 772–772.

Darwin C. 1877. On the various contrivances by which british and foreign orchids are fertilised by insects. New York: Appleton and CO.

Donoghue MJ, Sanderson MJ. 2015. Confluence, synnovation, and depauperons in plant diversification. The New phytologist 207: 260–74.

Donoghue MJ, Winkworth R. 2005. Viburnum phylogeny based on combined molecular data: implications for taxonomy and biogeography. American Journal of Botany 92: 653–666.

Drummond AJ, Bouckaert RR. 2014. Bayesian evolutionary analysis with BEAST 2. Cambridge University Press.

Drummond C, Eastwood R, Miotto S, Hughes CE. 2012. Multiple continental cadiations and correlates of diversification in Lupinus (Leguminosae): testing for key innovation with incomplete taxon sampling. Systematic Biology 61: 443–460.

Fitzjohn RG. 2012. Diversitree: comparative phylogenetic analyses of diversification in R. Methods in Ecology and Evolution 3: 1084–1092.

Gentry AH. 1982. Neotropical floristic diversity: phytogeographical connections between Central and South America, Pleistocene climatic fluctuations, or an accident of the Andean orogeny? Annals of the Missouri Botanical Garden 69: 557–593.

Gentry AH, Dodson CH. 1987. Diversity and biogeography of Neotropical vascular epiphytes. Annals of the Missouri Botanical Garden 74: 205–233.

Gerlach G. 2011. The genus Coryanthes: a paradigm in ecology. Lankesteriana 11: 253–264.

Gerlach G, Pérez-Escobar OA. 2014. Looking for missins swans: Phylogenetics of Cycnoches. Orchids 83: 434–437.

Gerlach G, Schill R. 1991. Composition of Orchid Scents Attracting Euglossine Bees. Botanica Acta 104: 379–384.

Givnish TJ, Spalink D, Ames M, Lyon SP, Hunter SJ, Zuluaga A, Doucette A, Caro GG, Mcdaniel J, Clements MA, et al. 2016. Orchid historical biogeography, diversification, Antarctica and the paradox of orchid dispersal. : 1–12.

Givnish TJ, Spalink D, Ames M, Lyon SP, Hunter Z, Zuluaga A, Iles W, Clements MA, Arroyo MT, Leebens-Mack J, et al. 2015. Orchid phylogenomics and multiple drivers of their extraordinary diversification. Proceedings of the Royal Society B: Biological Sciences 282: 20151553.

Goldberg E, Lancaster L, Ree RH. 2011. Phylogenetic inference of reciprocal effects between geographic range evolution and diversification. Systematic Biology 60: 451–465.

Gregory-Wodzicki KM. 2000. Uplift history of the Central and Northern Andes: a review. Geological Society of America Bulletin 112: 1091–1105.

von Hagen KB, Kadereit JW. 2003. The diversification of Halenia (Gentianaceae): ecological opportunity versus key innovation. Evolution 57: 2507–2518.

Heibl C. 2014. MEGAPTERA Mega-Phylogeny Techniques in R.: 1–7.

Hijmans RJ, Elith J. 2016. Species distribution modeling with R.

Hoorn C, Guerrero J, Sarmiento GA, Lorente MA. 1995. Andean tectonics as acause for changing drainage patterns in Miocene northern South America. Geology 23: 237–240.

Hoorn C, Mosbrugger V, Mulch A, Antonelli A. 2013. Biodiversity from mountain building. Nature Geoscience 6: 154–154.

Hoorn C, Wesselingh FP, Steege H, Bermudez M a, Mora A, Sevink J, Sanmartín I, Anderson CL, Figueiredo JP, Jaramillo C, et al. 2010. Amazonia Through Time: Andean uplift, climate change, landscape evolution, and biodiversity. Science 330: 927–931.

Hughes CE, Atchison GW. 2015. The ubiquity of alpine plant radiations: from the Andes to the Hengduan mountains. New Phytologist 207: 275–282.

Hughes C, Eastwood R. 2006. Island radiation on a continental scale: exceptional rates of plant diversification after uplift of the Andes. Proceedings of the National Academy of Sciences of the United States of America 103: 10334–10339.

Hughes CE, Pennington RT, Antonelli A. 2013. Neotropical plant evolution: assembling the big picture. Botanical Journal of the Linnean Society 171: 1–18.

Jost L. 2004. Explosive local radiation of the genus Teagueia (Orchidaceae) in the upper Pastaza watershed of Ecuador. Lyonia 7: 41–47.

Karremans AP. 2016. Genera Pleurothallidinarum: an updated phylogenetic overview of Pleurothallidinae. Lankesteriana 16: 219–241.

Kolanowska M. 2014. The orchid flora of the Colombian Department of Valle del Cauca. Revista Mexicana de Biodiversidad 85: 445–462.

Kreft H, Koster N, Kuper W, Nieder J, Barthlott W. 2004. Diversity and biogeography of vascular epiphytes in Western Amazonia, Yasuni, Ecuador. Journal of Biogeography 31: 1463–1476.

Krömer T, Gradstein SR. 2003. Species richness of vascular epiphytes in two primary forests and fallows in the Bolivian Andes. Selbyana 24: 190–195.

Küper W, Kreft H, Nieder J, Köster N, Barthlott W. 2004. Large-scale diversity patterns of vascular epiphytes in Neotropical montane rain forests. Journal of Biogeography 31: 1477–1487.

Lagomarsino L, Condamine FL, Antonelli A, Mulch A, Davis CC. 2016. The abiotic and biotic drivers of rapid diversification in Andean bellflowers (Campanulaceae). New Phytologist 210: 1430–1432.

Lindley J. 1843. Catasetidae. Edwards’s Botanical Register 29: sub t. 22

Luebert F, Hilger HH, Weigend M. 2011. Diversification in the Andes: age and origins of South American Heliotropium lineages (Heliotropiaceae, Boraginales). Molecular Phylogenetics and Evolution 61: 90–102.

Lugo MA, Ferrero M, Menoyo E, Estévez MC, Siñeriz F, Anton A. 2008. Arbuscular mycorrhizal fungi and rhizospheric bacteria diversity along an altitudinal gradient in South American Puna grassland. Microbial Ecology 55: 705–713.

Madriñán S, Cortés AJ, Richardson JE. 2013. Páramo is the world’s fastest evolving and coolest biodiversity hotspot. Frontiers in Genetics 4: 1–7.

Matzke NJ. 2013. Probabilistic historical biogeography: new models for founder-event speciation, imperfect detection, and fossil allow improved accuracy and model-testing. Frontiers of biogeography 5: 243–248.

Matzke NJ. 2014. Model selection in historical biogeography reveals that founder-event speciation is a crucial process in island clades. Systematic Biology 63: syu056-.

McGuire JA, Witt CC, Remsen J V, Corl A, Rabosky DL, Altshuler D, Dudley R. 2014. Molecular phylogenetics and the diversification of Hummingbirds. Current Biology 24: 1–7.

Merckx VSFT, Hendriks K, Beentjes K, Mennes C, Becking LE, Peijnenburg KTCA, Afendy A, Arumugam N, Boer H De, Biun A, et al. 2015. Evolution of endemism on a young tropical mountain. Nature 524: 347–350.

Miller MA, Schwartz T, Pickett BE, He S, Klem EB, Scheuermann RH, Passarotti M, Kaufman S, Leary MAO. 2015. A RESTful API for access to phylogenetic tools via the CIPRES Science Gateway. Evolutionary Bioinformatics 11: 43–48.

Morlon H, Lewitus E, Condamine FL, Manceau M, Clavel J, Drury J. 2016. RPANDA: An R package for macroevolutionary analyses on phylogenetic trees. Methods in Ecology and Evolution 7: 589–597.

Myers N, Mittermeier RA, Mittermeier CG, Fonseca GAB, Kent J. 2000. Biodiversity hotspots for conservation priorities. Nature 403: 853–858.

Nevado B, Atchison GW, Hughes CE, Filatov DA. 2016. Widespread adaptive evolution during repeated evolutionary radiations in New World lupins. Nature Communications 7: 1–9.

Pandey M, Sharma J, Taylor DL, Yadon VL. 2013. A narrowly endemic photosynthetic orchid is non-specific in its mycorrhizal associations. Molecular Ecology 22: 2341–2354.

Pansarin LM, Castro MDEM, Sazima M. 2009. Osmophore and elaiophores of Grobya amherstiae (Catasetinae, Orchidaceae) and their relation to pollination. Botanical Journal of the Linnean Society 159: 408–415.

Parra-Sánchez E, Retana J, Armenteras D. 2016. Edge Influence on diversity of orchids in Andean cloud forests. Forests 7: 1–13.

Pennington RT, Lavin M, Särkinen T, Lewis GP, Klitgaard BB, Hughes CE. 2010. Contrasting plant diversification histories within the Andean biodiversity hotspot. Proceedings of the National Academy of Sciences 107: 13783–13787.

Pérez-Escobar OA, Balbuena JA, Gottschling M. 2016a. Rumbling orchids : How to assess divergent evolution between chloroplast endosymbionts and the nuclear host. Systematic Biology 65: 51–65.

Pérez-Escobar OA, Gottschling M, Whitten WM, Salazar G, Gerlach G. 2016b. Sex and the Catasetinae (Darwin’s favourite orchids). Molecular Phylogenetics and Evolution 97: 1–10.

Pirie MD, Chatrou LW, Mols JB, Erkens RHJ, Oosterhof J. 2006. ‘Andean-centred’ genera in the short-branch clade of Annonaceae: Testing biogeographical hypotheses using phylogeny reconstruction and molecular dating. Journal of Biogeography 33: 31–46.

Pridgeon AM, Cribb PJ, Chase MW, Rasmussen FN. 2009. Genera Orchidacearum: Epidendroideae (part two), (Eds.). Vol. 5. Oxford: Oxford University Press.

Pupulin F, Karremans AP, Gravendeel B. 2012. A reconsideration of the empusellous species of Specklinia (Orchidaceae: Pleurothallidinae) in Costa Rica. Phytotaxa 63: 1–20.

Rabosky DL. 2014. Automatic detection of key innovations, rate shifts, and diversity-dependence on phylogenetic trees. PLoS ONE 9.

Rabosky DL, Donnellan SC, Grundler M, Lovette IJ. 2014. Analysis and visualization of complex Macroevolutionary dynamics: An example from Australian Scincid lizards. Systematic Biology 63: 610–627.

Ramirez SR, Eltz T, Fujiwara MK, Gerlach G, Goldman-Huertas B, Tsutsui ND, Pierce NE. 2011. Asynchronous Diversification in a Specialized Plant-Pollinator Mutualism. Science 333: 1742–1746.

Ramírez SR, Gravendeel B, Singer RB, Marshall CR, Pierce NE. 2007. Dating the origin of the Orchidaceae from a fossil orchid with its pollinator. Nature 448: 1042–1045.

Ramsey J, Ripley B. 2010. pspline: penalized smoothing splines

Richter M, Diertl K, Emck P, Peters T, Beck E. 2009. Reasons for an outstanding plant diversity in the tropical Andes of Southern Ecuador. Landscape Online 12: 1–35.

Santos JC, Coloma L a., Summers K, Caldwell JP, Ree R, Cannatella DC. 2009. Amazonian amphibian diversity is primarily derived from late Miocene Andean lineages. PLoS Biology 7: 0448–0461.

Scott A, Ram K, Hart T, Chamberlain MS. 2015. Package ‘ spocc ’.

Simpson GG. 1953. The major features of evolution. New York: Columbia University Press.

Singer RB. 2002. The pollination mechanism in Trigonidium obtusum Lindl (Orchidaceae: Maxillariinae): sexual mimicry and trap-flowers. Annals of Botany 89: 157–163.

Smith BT, McCormack JE, Cuervo M, Hickerson MJ, Aleixo A, Burney CW, Xie X, Harvey MG, Faircloth BC, Cadena CD, et al. 2014. The drivers of tropical speciation. Nature 515: 406–409.

Stamatakis A. 2014. RAxML version 8: A tool for phylogenetic analysis and post-analysis of large phylogenies. Bioinformatics 30: 1312–1313.

Ter Steege H, Pitman NC, Sabatier D, Baraloto C, Salomão RP, Guevara JE, Phillips OL, Castilho C V, Magnusson WE, Molino J-F, et al. 2013. Hyperdominance in the Amazonian tree flora. Science 342: 325–342.

Talavera G, Castresana J. 2007. Improvement of phylogenies after removing divergent and ambiguously aligned blocks from protein sequence alignments. Systematic Biology 56: 564–577.

Tremblay RL, Ackerman JD. 2001. Gene flow and effective population size in Lepanthes (Orchidaceae): a case for genetic drift. Biological Journal of the Linnean Society 72: 47–62.

Uribe-Convers S, Tank DC. 2015. Shifts in diversification rates linked to biogeographic movement into new areas: An example of a recent radiation in the Andes. American Journal of Botany 102: 1–16.

Valencia R, Balslev H, Miño C G. 1994. High tree alpha-diversity in Amazonian Ecuador. Biodiversity and Conservation 3: 21–28.

Vargas P, Zardoya R. 2014. The tree of life. Sunderland: Sinauer Associates.

Verboom GA, Archibald JK, Bakker FT, Bellstedt DU, Conrad F, Dreyer L, Forest F, Galley C, Goldblatt P, Henning J, et al. 2009. Origin and diversification of the Greater Cape flora: Ancient species repository, hot-bed of recent radiation, or both? Molecular Phylogenetics and Evolution 51: 44–53.

Whitten WM, Neubig KM, Williams NH. 2014. Generic and subtribal relationships in neotropical Cymbidieae (Orchidaceae) based on matK/ycf1 plastid data. Lankesteriana 13: 375–392.

Willis JC. 1922. Age and area. London: Cambridge University Press.

Zotz G, Winkler U. 2013. Aerial roots of epiphytic orchids: the velamen radicum and its role in water and nutrient uptake.: 733–741.

